# Cognitive Control Adjusts to Behavioral Constraints at Short Timescales

**DOI:** 10.64898/2026.06.12.731633

**Authors:** Camila Losada, Arno Feinstein, Alexis Monnet-Aimard, Guilhem Ibos

**Author notes:** Each author contributed equally to this work (Camila Losada, Arno Feinstein, Alexis Monnet-Aimard).

## Abstract

Cognitive functions encompass a large set of abstract constructs used for adapting our behavior to environmental constraints, each of them acting at specific timescales. For example, while working memory operates over several seconds or minutes, decision making occurs over much shorter periods. Solving a behavioral task thus relies on specific cognitive strategies that use several functions over time. Here, we investigated how two macaque monkeys coordinate working memory, selective attention, decision-making and executive control of eye movements during performance of a modified delay match-to-sample task. The economy of this task (including reward expected value and cost of errors) evolves on short timescales within trials. In addition, this task allowed us to manipulate engagement of cognitive resources at longer timescales. Using tools from signal detection theory, we closely analyze how monkeys’ performances evolve over time and infer their specific strategies in terms of control of cognitive functions. In addition, during covert attention, fixational eye movements and pupil size, provided reliable markers of slow variations in cognitive state across trials, although not of rapid within-trial changes in cognitive control. Together, these results show that each monkey adapted to the same task by implementing individual, dynamically evolving cognitive strategies across multiple timescales.

## Introduction

Adapting our behavior to our environment requires developing task-specific behavioral strategies which often coordinate several cognitive, sensory and motor functions. In order to maximize success during our daily tasks, we simultaneously hold on to our goals, orient sensory processing, integrate relevant information, avoid distraction while staying alert to relevant contextual switches and exert precise control over our motor plan. Importantly, the different timescales (Janssens et al., 2016; Aben et al., 2017; Soltani et al., 2021) at which cognitive constructs (Uddin, 2021) and task contingencies operate imply intricately coordinating each of these processes. However, it is unclear how such coordinated behavioral strategies emerge and adapt on short timescales.

For instance, *working memory* stores, manipulates and protects information over the course of seconds or minutes (Baddeley, 1992; Goldman-Rakic, 1995; Pasternak and Greenlee, 2005). *Top-down selective attention* (Moran and Desimone, 1985; Desimone and Duncan, 1995; Connor et al., 2004), gates the bottom-up flow of sensory information for better downstream integration (Ibos and Freedman, 2014, 2016) and can be adjusted quickly. In parallel, different task contingencies may impose specific constraints on executive functions such as motor preparation, control of eye movements, or timely execution of motor plans. As a result, in order to generate appropriate motor plans, *decision-making* processes requires to integrate sensory and working memory information (Gold and Shadlen, 2007; Freedman and Ibos, 2018). In the context of dynamic tasks (Brehmer, 1992), decision making benefits from updating representations of quickly evolving reward utility (Loewenstein et al., 2008; Botvinick and Braver, 2015; Westbrook and Braver, 2015), including the expected value associated with successful behaviors (reward value given the effort required to receive it) and costs associated with different types of errors (Kable and Glimcher, 2009; Padoa-Schioppa, 2011; Westbrook et al., 2013). Coordination of all these cognitive processes result in specific decision strategies which may therefore change at different timescales depending on the specific economical constraints of behavioral tasks.

In this study, we explore how behavioral strategies of two macaque monkeys performing a task designed for future electrophysiological investigation evolve within each trial over short timescales. Importantly, the utility of different types of behavioral responses evolved within each trial in a predictable manner, allowing animals to adjust their strategies accordingly. Both monkeys performed a modified version of a visual delayed match to sample task. We show that both monkeys adopted different decision strategies based on task contingencies. We link strength of working memory encoding to stimulus discriminability and decision criterion. Finally, we show that during covert attention, fixational eye movements and pupil size provided reliable markers of slow variations in cognitive state across trials, although not of rapid within-trial changes in cognitive control. Together, these results show that each monkey adapted to the same task by implementing individual, dynamically evolving cognitive strategies across multiple timescales.

## Results

### Behavioral task

We trained two animals (monkey R and monkey M) to perform a delayed conjunction-matching task (Figure 1). Monkeys held a manual lever located inside the chair they were sitting in. A fixation point appeared at the center of the screen. They had to maintain gaze on this point until trials end (within a 2° radius virtual window). Any fixation breaks resulted in immediate trial interruption. A sample stimulus was presented at one of two locations (left or right) for 450 ms. After a variable delay, a succession of one to four test stimuli was presented at the same location as the sample stimulus, simultaneously with as many distractor stimuli located on the opposite hemifield. Test and distractor stimuli were a conjunction of one of eight equally spaced orientations and one of eight equally spaced colors (Figure 1B). Sample stimuli were a conjunction of one of two orientations (orientation 1: o1, or orientation 5: o5) and one of two colors (color 1: c1 or color 5: c5). For the rest of the article, each of the four sample stimuli are referred to according to their orientation and color as o1c1, o1c5, o5c1 and o5c5. In order to receive a reward, monkeys had to release the lever when a test stimulus matched sample stimulus’ location, orientation and color. In addition, sample stimuli could also be composed of a neutral color (central to test color space) and a neutral orientation (non-oriented dot). No target stimuli were ever presented during both neutral (predictable from sample onset) and catch trials (unpredictable until the last test stimulus). In these trials, monkeys were rewarded for holding the lever and fixation until the end of the trial.

**Figure 1:**
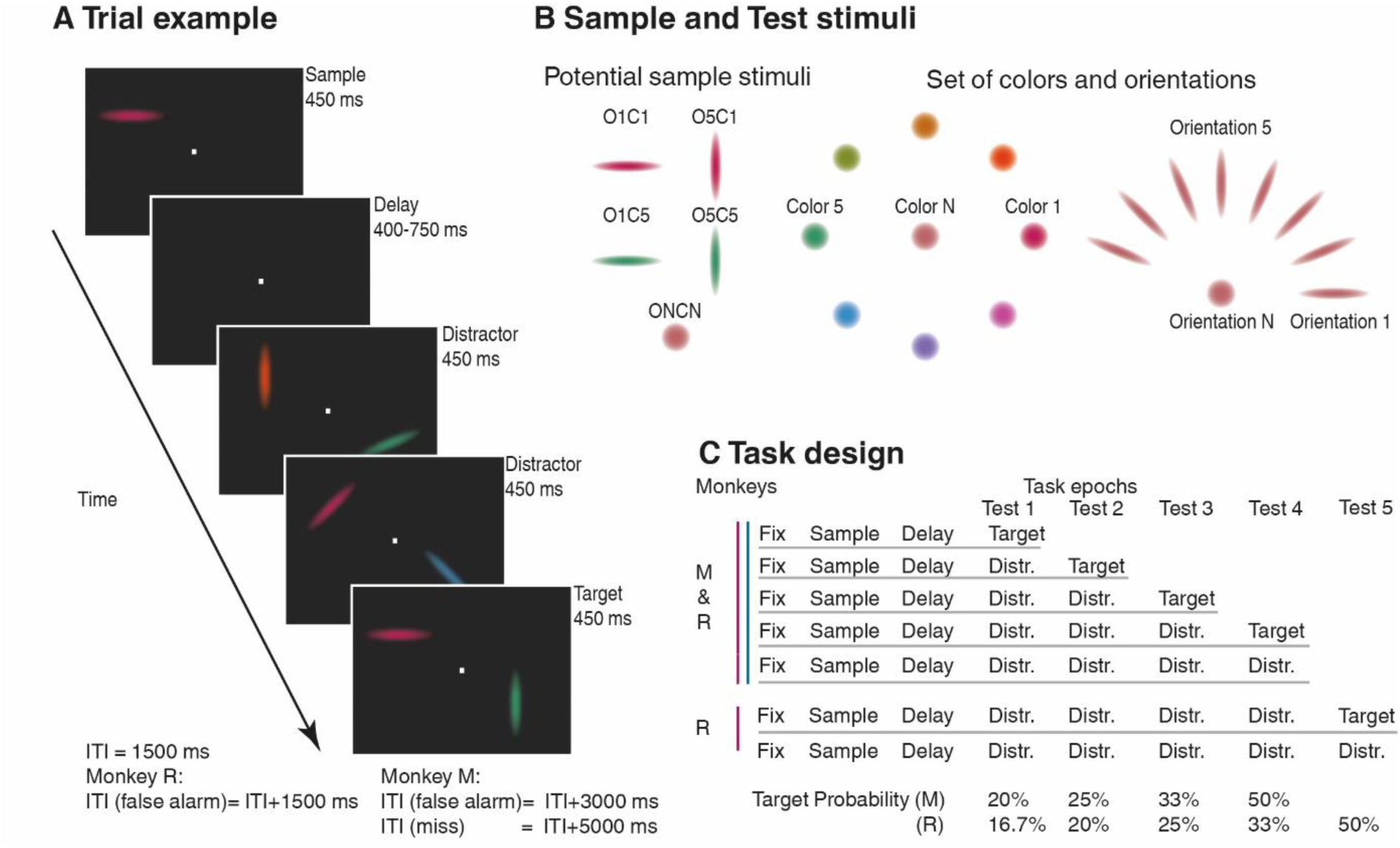
Delayed conjunction matching task. A. Example of trial: sample stimulus (o1c1) is presented on one of two locations (left or right), followed by a delay and a succession of test stimuli. Monkeys (M and R) must match the location, color and orientation of test and sample stimuli. B. Set of visual features for sample and test (peripheral) stimuli. C. Task design for each monkey (M and R) with target probability.

Task design (Figure 1C) differed slightly for both monkeys. In monkey R, the initial design used during the first 32 sessions included five possible test epochs. As trial progressed, probability that a target would appear increased across epoch, from 16.5% (1/6) at test epoch #1 (T1), 20 % (1/5) at test epoch #2 (T2), 25% (1/4) at test epoch #3, 33% (1/3) at test epoch #4 (T4) to 50% (1/2) at test epoch #5 (T5). To speed up trials and increase motivation, one test epoch was removed in the remaining 22 sessions, resulting in four possible test epochs and a corresponding increase in target probability from 20% at T1 to 50% at T4. The same four test epoch design was used across all monkey M’s 24 sessions.

### General performances

We first assessed monkeys’ general performances for each target stimuli (Figure 2A). We classified trials as hits, false alarms or misses. Both monkeys showed different performance profiles. Monkey R showed a high proportion of correct responses across sample identity, with ∼80 % hit trials, ∼10% false alarm trials and ∼10% miss trials. Monkey M showed a stronger tendency to miss target stimuli, with ∼60% hit trials, ∼10% false alarm trials and ∼30% miss trials. Interestingly, both monkeys performed similar amount of correct trials for each session (monkey R: 516 +/- 152 trials/session; monkey M: 535 +/- 134 trials/session, t-test, p=0.6), however monkey M sessions (mean 130.2 +/- 17 minutes/session) lasted significantly longer than monkey R’s (mean 89 minutes +/-17 minutes/session; t-test, p<0.001). Such effect is fully explained by the differences of cumulative penalties applied to each monkey’s error trials (mean 48.15 minutes/session). Higher miss rates may reflect different processes which will be explored in the following sections. For example, they could be explained by a more selective decision threshold in monkey M compared with monkey R. Another explanation could be related to lower motivation or weaker cognitive control, such as a reduced ability to keep task-relevant information in working memory over the course of trials and to orient attention toward relevant representations. The following analyses aim at identifying monkey R’s and M’s respective strategies.

**Figure 2:**
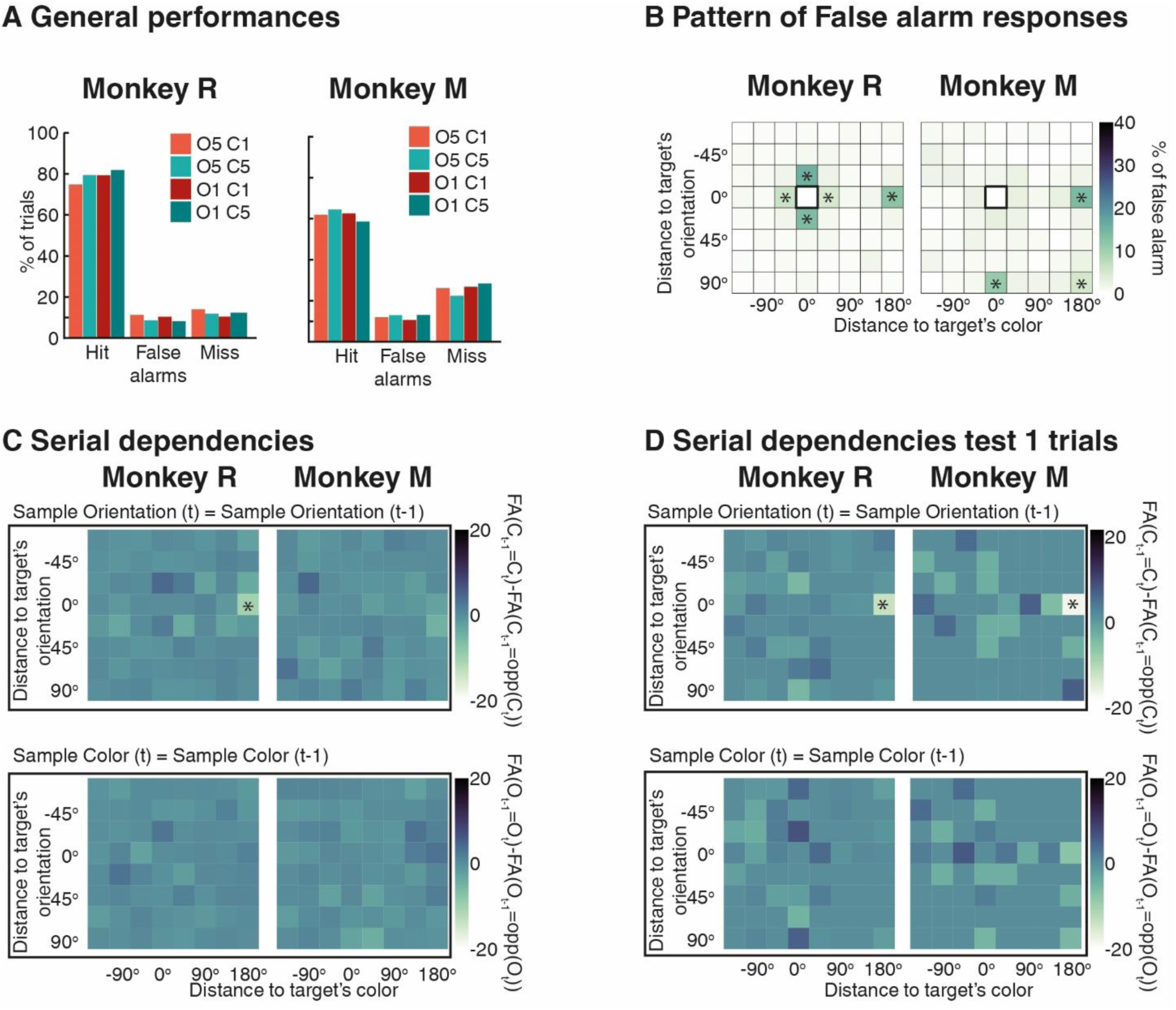
A. Behavioral performances: Proportions of hit, false alarm and miss trials for each sample and both monkeys. B. Percentage of false alarm responses for each test stimulus as a function of orientation and color distance to current sample stimuli. C. Serial dependencies between previous and current sample stimuli. Top, samples’ orientation of previous and current trials match (testing color serial dependency). Bottom, samples’ color of previous and current trials match (test orientation serial dependency). For each panel, each line and column correspond to the orientation and color (respectively) of currently presented test stimuli. Color code corresponds to differences of percentage of false alarm responses between trials during which previous sample matched current sample’s color (top) or orientation (bottom) with the ones during which previous sample didn’t match current sample’s color (top) or orientation (bottom).(*, Bonferonni corrected t-test, p<0.01). D. Serial dependencies between previous and current trials (T1 only). Similar to C, instead (*, fdr corrected t-test, p<0.01)

### False alarm patterns

False alarm responses can inform us about how attentional and memory resources are allocated during trials and may reveal specific search strategies for each monkey. We therefore analyzed false alarm rates for each of the 64 potential test stimuli. To do so, we pooled trials across sample identities and reorganized test stimuli according to their distance from sample (and target) in color and orientation space (Figure 2B). Both monkeys exhibited markedly different false alarm patterns.

Monkey R made false alarms mainly to two types of stimuli. First, false alarms were more frequent for stimuli that were highly similar to current target in both orientation and color (± 27.5° from target’s orientation, ± 45° from target’s color). This error pattern may reflect the perceptual complexity of the task, with orientation discrimination threshold close to 27.5° and color discrimination threshold close to 45°. Interestingly, stimuli sharing only one target’s feature (orientation or color) triggered false alarm. For example, stimuli composed of a color (or an orientation) adjacent to target triggered false alarm only when they matched the target orientation (or color respectively). This pattern suggests that monkey R’s target detection strategy results, to some extent, on the joint evaluation of bound color and orientation information rather than on either feature alone. Second, monkey R also showed high false alarm rates for stimuli composed of target’s orientation combined with the color opposite to the target color in the stimulus set (180° away from it). These stimuli shared the target orientation but were colored like another potential target (o1c5 test stimuli during o1c1 sample trials, or o5c1 test stimuli during o5c5 sample trials). Because previous studies have shown that behavior can be biased by information kept in working memory on previous trials, a phenomenon described as serial dependence (Barbosa and Compte, 2020; Barbosa et al., 2020; Cicchini et al., 2024) we tested the relationship between these false alarm errors and previous trial’s relevant color. We therefore computed false alarm rates as a function of the distance between the current and previous sample stimuli (Figure 2C). This analysis shows that false alarms to orientation-matching associated with other colors were more likely when that color had been relevant on the previous trials, consistent with a serial dependence effect on color. By contrast, the orientation of the previous sample did not appear to influence the current false alarm pattern. Together, these results suggest that, in monkey R, the representation of the task-relevant color was partially biased by the previous trial.

Monkey M showed a drastically different pattern of false alarm (Figure 2B). Unlike monkey R, he correctly rejected stimuli that were similar to target, suggesting sharper perceptual discrimination or, at least, a stricter response criterion for near-target stimuli. Instead, he more often responded to other potential targets (e.g. o1c5, o5c1 or o5c5 during o1c1 trials). Such errors could potentially reflect weaker maintenance of sample identity in working memory, or reduced inhibitory control over responses to any of these potential targets. However, this error pattern was not explained by serial dependence from the previous trial, as monkey M did not show a clear dependence for neither the color nor the orientation of the previous sample stimuli (Figure 2C). Since serial dependence is thought to correlate with the strength of relevant information encoding in working memory during previous trial, lack of serial dependence over trials is therefore consistent with weak encoding of information in working memory. Taken together, monkey’s M high miss rate (Figure 2A) and false alarm pattern (Figure 2B) suggest that his behavior was less strongly guided by stable representations of current sample stimuli.

### Evolution of target probability and cognitive control over time

To better characterize each monkey’s behavioral strategy, we evaluated performances at each test epoch. The structure of the task allows us to analyze behavior within the general framework of signal detection theory (SDT, Figure 3 top). At each test epoch, monkeys had to detect and report a matching stimulus. In addition, they were required to maintain fixation along trials. They therefore faced two main constraints: first, detecting and reporting target stimuli while avoiding responses to distractor stimuli, and second, maintaining gaze fixation within a 2° radius virtual window. Reporting a target was categorized as a hit, whereas failing to report it was categorized as a miss. Similarly, responding to or ignoring distractor stimuli were respectively categorized as false alarms or correct rejections. In such framework, performance depends on two factors: target/distractor discriminability (sensitivity), captured by d’ measurement; and a threshold (criterion), above which an observer would make a response. For a fixed criterion, reducing d’ would result in decreasing hit rates: *hit / (hit + miss),* and increasing false alarm rates: *false alarm / (false alarm + correct rejections)*. Inversely, increasing d’ would result in increasing hit rates and decreasing false alarm rates. Conversely, for a fixed level of sensitivity, decreasing criterion would result in increasing both hit and false alarm rates, whereas increasing criterion would result in decreasing both.

**Figure 3:**
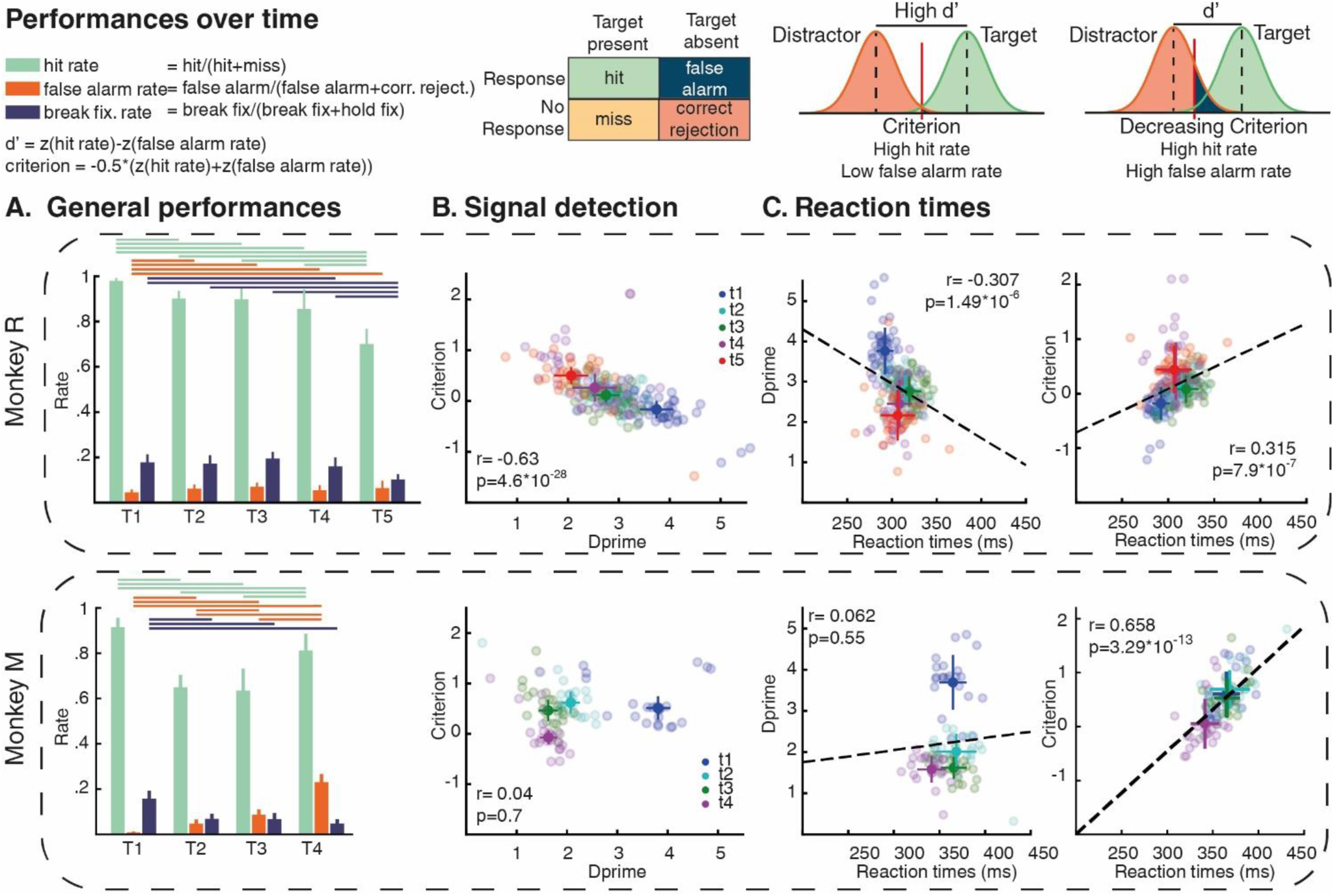
Performances over time. Top: Principles of signal detection theory. A. Evolution over each test epoch of median (± absolute deviation) hit-rate (green), false alarm-rate (orange), and fixation-break (blue) rate for monkey R (top) and monkey M (bottom) (horizontal bars, p<0.01, repeated measure ANOVA). B: Evolution at each test epoch of criterion and d’ for each test session (transparent dots), full dots and bars are means ± std. C. Correlation between each session’s averaged reaction time with d’(left) and criterion (right), (dotted black line: linear regression, r: pearson corr coef, T-test).

Moreover, it is important to highlight that each trial consisted of a succession of one to four (five for some of monkey R sessions) test stimuli (Figure 1C). As trials progressed, the probability that a target would appear increased at each test epoch. Simultaneously, monkeys had already invested more time and effort in maintaining fixation, as each additional test epoch extended trial duration by 450 ms. Thus, as trials unfolded, both the probabilistic structure of the task and its cumulative behavioral cost changed. This structure allows us to draw specific hypothesis about potential detection and response policies strategies which can be directly addressed with SDT tools. Larger efforts (associated to longer fixation and search) result in decreasing the expected value for reward associated to correct trials (hit and correct rejection during catch trials) and increasing the cost associated to error trials (miss and false alarm trials as well as break-fixation). One possible strategy would therefore be to have a less stringent response policy (for example by reducing executive control over detection-decision or manual response) to maximize target detection. This strategy would result in increasing hit rates and false alarm rates (decreasing criterion). However, because reward amount was similar across trials, cost associated to errors also increase with trial duration, prompting animals to reduce errors unrelated to target detection, such as fixation-break. In parallel, as trials unfolded, probability of catch trials also increased at each test epoch. Under these conditions, another potential strategy would be to reduce risk and behave as if the current trial were increasingly likely to be a catch trial (no target presented, no response needed). To determine which strategy each animal developed, we computed hit rates, false alarm rates and fixation-break rates for each test epoch (Figure 3A).

Monkey R’s hit rates, and fixation-break rates decreased across test epochs (ANOVA with repeated measure, p < 0.01). In addition, false alarm rates slightly increased (from ∼4% for test epoch #1 to 7% for test epoch #5, ANOVA with repeated measure, p < 0.01). Based on hit and false alarm rates, we computed d’ and criterion for each test epoch (Figure 3B). Monkey R’s performance slightly drifted over the course of the trial. Performance was best at the first test epoch, with high d’ and low criterion. Across successive test epochs, d’ decreased while criterion increased (Pearson correlation, p < 0.01). This pattern suggests a progressive reduction in discriminability (supposedly linked to decreased attention) and an increased criterion (supposedly linked to stricter response policy). In addition, fixation-break rates decreased across test epoch, suggesting tighter control over eye movements to prevent errors. Overall, monkey R became more conservative at the end of each trial, limiting both manual responses and fixation-break errors, in a manner consistent with an increasing tendency to bet on catch trial.

Monkey M’s performances were, once again, dramatically different and varied rapidly along trials. During the first test epoch, hit rates were very high and false alarms rates were very low, corresponding to high d’ and high criterion (Figure 3A and B). This pattern indicates that monkey M was initially able to discriminate test stimuli’s color and orientation and correctly kept information in working memory over the delay period. However, hit rates dropped to ∼60% at the following test epochs (2 and 3), while false alarm rates increased to ∼10%, consistent with relatively low d’ (∼2) and high criterion. Such drop-out in performance may reflect a reduction in cognitive control, for example through weaker strength of stimuli kept in working memory or attention paid to test stimuli’s feature, independently of response policies modulation. At the fourth test epoch, monkey M showed increased hit rates coupled with increased false alarm rates, corresponding to lower discriminability (d’ < 2) and low criterion. Monkey M is therefore prompt to execute a manual response at the final test epoch. Together with the pattern of false alarm responses, these results suggest that monkey M bet on match stimuli at the final test epoch. The relatively sharp decrease of d’ between the first and subsequent test epochs, together with high false alarm rate for potential targets as well as the lack of serial dependence between successive trials, are consistent with decreased strength of attention and working memory over time. If this hypothesis is correct and if monkey M properly used information kept in working memory more effectively at the first test epoch before this cognitive control weakened, we should observe serial dependence either when false alarms occur during the first test epoch (strong memory encoding), or when previous trial’s target stimuli occurred during the first test epoch. As shown in Figure 2D, we observed a strong increase of previous trial’s color on false alarm rates at the first test epoch, an effect that was particularly striking in monkey M. Unfortunately, we did not have enough false alarm trials in which target of the previous trials was presented at the first test epoch to test the second prediction reliably. Together, these results support our hypothesis that monkey M rapid changes in cognitive state after the first test epoch and again following the third test epoch, reflect dynamic adjustments in cognitive control, especially in the encoding strength of information in working memory and in decision criterion.

### Discriminability, criterion and reaction times

We next explored how behavioral strategies, expressed in terms of d’ and criterion, related to changes in reaction times (Figure 3C). It is important to note that behavioral responses were constrained by the 450 ms of target presentation, putting strong time pressure on both decision-making and motor processes.

Despite these constraints, monkeys R and M showed distinct dynamics in reaction times. In monkey R, reaction times at test epoch 1 (mean: 292 ± 9 ms) increased slightly at test epochs 2 (315 ± 12) and 3 (319 ± 14 ms,) before decreasing at test epochs 4 (308 ± 14 ms) and 5 (306 ± 19 ms) (ANOVA for repeated measures, p < 0.01). In monkey R, whose d’ and criterion varied in a correlated manner, variations in reaction times were significantly correlated with both d’ (Pearson correlation, r = -0.307, p < 0.01) and criterion (Pearson correlation, r = 0.315, p < 0.01).

In Monkey M, reaction times were stable and high during test epochs 1 (365 ± 15 ms), 2 (369 ± 22 ms) and 3 (366 ± 14 ms), but decreased at test epoch 4 (341 ± 16 ms) (ANOVA for repeated measures, p < 0.01). Interestingly, monkey M’s reaction times were not significantly correlated with d’ (Pearson correlation, r = 0.062, p = 0.55), but were strongly correlated with criterion (Pearson correlation, r = 0.658, p < 0.01).

Together, these results provide insight into how both monkeys’ strategies and cognitive states evolved over the course of the trial, suggesting that the control of cognitive functions can affect behavior on relatively short timescales.

### Cognitive state, eye movements and pupil size

Cognitive states related to different levels of cognitive control (such as maintaining information in working memory or deploying covert attention) are known to strongly influence small eye movements (Engbert and Kliegl, 2003; Martinez-Conde et al., 2013; Rosen and Freedman, 2025). We therefore tested whether monkeys’ cognitive states could be inferred from gaze position within the virtual fixation window, from micro-saccade rate and from variations in pupil-size. In this section, we specifically compared trials in which the sample stimulus was non-neutral (o1c1, o1c5, o5c1 and o5c5), and therefore required greater cognitive effort, with trials in which the sample stimulus was neutral (oNcN), and therefore required minimal cognitive effort. We first analyzed gaze position within the fixation window during neutral and non-neutral trials as a function of test epochs (Figure 4). To do so, we computed the distance between actual gaze position and the projection, onto the fixation window, of the location of the relevant stimulus (Figure 4A). We used the average distance measured during a 200 ms beginning 50 ms following test stimulus onset (different time windows give similar results). In both monkeys (Figure 4D), gaze remained stable during neutral trials and was centered around the fixation point (distribution around 2°), and no effect of test epoch. By contrast, during non-neutral trials, gaze shifted closer to the border of the fixation window during the first test epoch (ANOVA, p < 0.001). From test epoch 2 to 4, gaze position went closer to central fixation point, with an average distance of around 2° in both monkeys.

**Figure 4:**
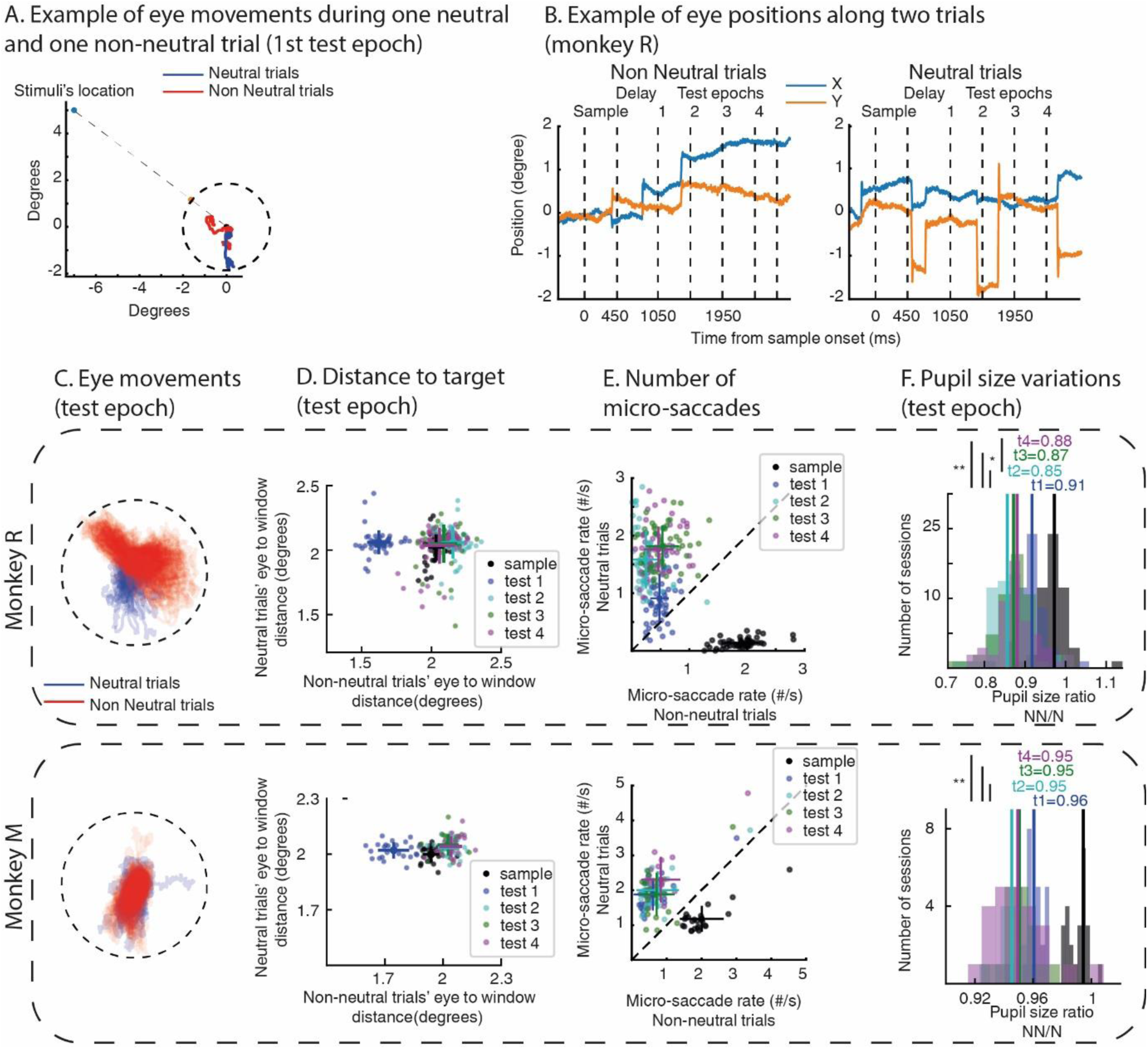
Eye movements and pupil size during sample/dealy period and test epochs in neutral and non-neutral trials. A. Example of eye movements during neutral (red) and non-neutral trials (blue). Blue dot represents stimulus location, orange dot represents its projection on fixation window. B. Example of eye coordinates during a neutral and a non-neutral trial. C. Overlayed eye position during a session for monkey R (top) and monkey M (bottom) during neutral (blue) and non-neutral trials (red). D Averaged distances between eye position and projection point during each epoch, for each session and each monkey (transparent dots). Full dots represent averaged position (+/- std). E. Micro-saccade rates during each trial epoch during neutral and non-neutral trials. Transparent dots represent each session, full dots represent averaged over session (+/-std). F. Pupil size ratio non-neutral / neutral for each task epoch (*, p<0.05; **, p<0.01).

We next tested oculomotor control by computing micro-saccade rate during each task epoch (sample and delay combined together, test epoch 1 to test epoch 4) for both neutral and non-neutral trials (Figure 4E). Micro-saccade rates showed a double dissociation between trial type and task epoch. During neutral trials, monkeys made few micro-saccades during sample and delay period. Micro-saccade rate increased sharply during subsequent test presentations. Inversely, during non-neutral trials, monkeys made more micro-saccade during sample and delay period, when sample information had to be maintained in working memory, but micro-saccade rate strongly decreased during test epochs, when attention was covertly directed toward the relevant location and features.

In addition, we compared pupil size between neutral and non-neutral trials at each test epoch (Figure 4F). We computed an index measuring the normalized ratio of pupil size in neutral versus non-neutral trials. For both monkeys, this index was systematically less than 1 (corresponding to more dilated pupils during non-neutral trials than during neutral trials) at each test epoch. However, we did not observe clear dynamics of pupil size across test epochs.

Finally, in order to link cognitive state to control of eye movements, we trained SVM classifiers to decode trial type (neutral versus non-neutral) over time for each session using either gaze position alone or gaze position combined with pupil size (Figure 5A). Including pupil size in the analyses strongly improved decoding performance for monkey M. Decoding accuracy strongly increased during the delay period and remained high across successive test epochs. In addition, we tested whether test epoch (1, 2, 3 or 4) could be decoded based on eye positions and pupil size for each session. Decoding accuracies (Figure 5B) based on gaze position alone were significantly above chance level (permutation test, p>0.05) for most of monkey’s R sessions during both non-neutral (N=47/51) and neutral trials (N=36/51). Including pupil size did not impact decoding accuracies significantly (permutation test, p>0.05). Monkey M eye-movements reflected test epochs only during neutral trials. We managed to decode test epoch during neutral trials only of a few sessions, using either position alone (N=8/24, permutation test, p<0.05) or in conjunction with pupil size (N=10/24, permutation test, p<0.05). Similar accuracies for neutral and non neutral trials in both animals suggest that oculomotor bio-markers reflect non-cognitive variables at this timescale such as tracking time within trials or before reward delivery.

**Figure 5.**
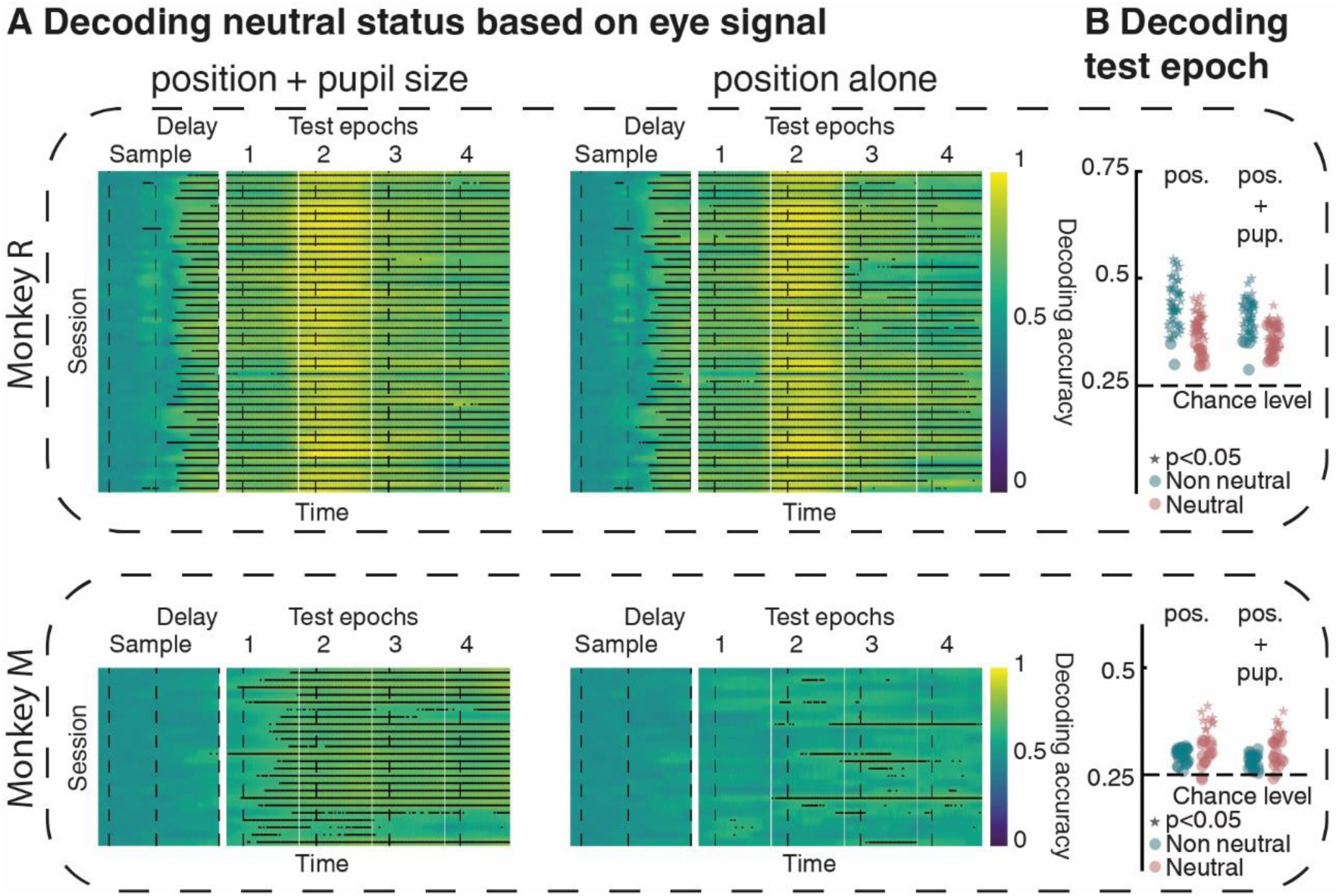
: A. Decoding trial type (neutral vs non neutral) based on eye position and pupil size (left) or eye position alone (right). Each line represents time course of classifier performances (color coded) for each session (*<0.05). B. Decoding test epoch (1 to 4) for each sessions’ eye position alone (left) or position and pupil size (right), during non-neutral (green) or neutral (brown) trials (permutation test, *<0.05).

Such results show that monkeys’ oculomotor behaviors, including pupil size were affected by slow cognitive processes taking place at long timescales (such as cognitive control during neutral vs non neutral trials), but did not vary with fast adjustment of search strategies taking place at short timescales (such as modulations of d’ and criterion at each test epoch).

## Discussion

In this study, we analyzed behavioral performances of two monkeys engaged in a task involving several cognitive processes: (i) maintaining the color, orientation, and location of sample stimulus in memory across the trial, (ii) attending to successive test stimuli’ visual features, (iii) evaluating whether these stimuli matched the sample, and (iv) executing the appropriate motor response while maintaining fixation. In parallel, the statistical structure of each trial evolved over time, potentially shaping the strategies developed by each animal on a short timescale.

We show that, although both monkeys performed the task with relatively high accuracy, they developed drastically different strategies across trial epoch. One monkey, monkey R, exhibited sustained cognitive control during trials, with marginal strategic drifts toward more conservative behavior. Interestingly, this monkey’s false alarm patterns reflected two distinct processes. First, color and orientation seemed to be computed separately and then bound together for decision-making. Second, the relevant color from the previous trial appeared to contaminate the current trial performance, consistent with a serial dependence-like effect.

The second monkey, monkey M, showed a drastically different performance pattern. This monkey exhibited very high performances during the first test epoch of the task, but d’ decreased sharply at following test epochs, a pattern consistent with reduced top-down cognitive control. In addition, the absence of serial dependence effect after the first test epoch may be associated with weaker encoding of information in working memory. Together, these interpretations are consistent with a decrease in cognitive control over both selective attention and working memory (Oberauer, 2019; Zhou et al., 2022; Badre, 2025). After the first test epoch, the sample representation kept in working memory may have weakened, leading to more automatic response to stimuli that were more likely to be targets (o1c1, o1c5, o5c1, o5c5). This is consistent with the observed decrease in decision criterion and faster reaction times at test epoch 4.

It is important to point out that such behavioral strategies, at least for test epoch 4, may also have been influenced by the penalty policy applied to monkey M’s error trials in order to increase his global detection rates. Because during training, this animal initially tended to respond predominantly to the first test epoch and to miss targets presented during subsequent test epochs, miss trials were followed by a strong penalty, with the subsequent intertrial interval increased to 6.5 s instead of 1.5 s. This manipulation was introduced with the goal of increasing his cognitive control. Under these conditions, monkey M may have adopted a lower decision threshold in order to avoid costly misses, thereby increasing both hit and false alarm rates independently of stimulus discriminability of working memory strength.

In this study, we also highlighted a tendency in monkey R (and to some extent monkey M) to make false alarm responses to stimuli composed of the relevant orientation and an irrelevant color. Such false alarms could reflect two non-mutually exclusive phenomena. First, monkey R may have adopted a search strategy primarily focused on detecting the relevant orientation, resulting in a higher false alarm rate for stimuli containing an alternative task-relevant color. However, such a strategy would result in specific false alarm patterns for stimuli surrounding the current target, with higher false alarm rates along the 0° orientation line than along the 0° color column. Our results do not support this prediction, as false alarm rates were higher for stimuli sharing the target’s color than for stimuli sharing the target’s orientation. Second, this effect could be related to contamination from previous trial, similar to a serial dependence effect (Barbosa et al., 2020; Houborg et al., 2023; Manassi et al., 2023; Cicchini et al., 2024). Our analyses suggest previous trial’s relevant color influenced current trial’s color decisions, and that this effect was linked to cognitive control. This effect does not completely correspond to how serial dependence has been described in previous studies. In those studies, the representation of a currently remembered feature is slightly biased toward the value of the corresponding feature remembered on the previous trial. These studies typically used graded response-scales, which are well suited to detecting serial dependence. Here, by contrast, we used a binary response (target is present or not). The type of serial dependence we describe therefore corresponds to an erroneous binding of features (relevant orientation with an irrelevant color that could be relevant in another trial context). Given this binary classification of response, being able to highlight such serial contamination at the behavioral level is particularly striking. Even more surprisingly, orientation representations seemed not to be affected by sample history, raising further questions about monkeys’ strategies used to bind visual features together. Several previous studies have used conjunctive stimuli in working memory task (Mante et al., 2013; Ibos and Freedman, 2014, 2016), and reanalyzing these existing datasets could inform us about how cognitive control impacts serial dependence under feature-binding demands.

Finally, recent studies have linked cognitive states to physiological signals such as pupil size (Kahneman and Beatty, 1966; Eckstein et al., 2017; Koevoet et al., 2026), facial (Tlaie et al., 2025), or eye movements (Rosen and Freedman, 2025). In the present study, we recorded only gaze position (constrained within a 2° radius virtual fixation window) and pupil size. Both variables differed reliably between neutral and non-neutral trials, reflecting broad differences in cognitive states. We were even able to decode trials’ neutral status within individual sessions using these measurements. However, these oculomotor and pupillary signals did not correlate with finer-grained indices of cognitive control such as d’ or criterion. One possible explanation is that these physiological variables and behavioral control metrics operate on partially different intrinsic timescales, especially considering the slow dynamics of pupil size (Mathôt and Vilotijević, 2023). Additional measures, such as heart rate, facial expressions or idiosyncratic body movements, including grip force on the manual lever, may provide a more sensitive readout of ongoing cognitive state.

## Methods

### Subjects and set up

All procedures conformed to European Commission law (DIRECTIVE 2010/63/EU OF THE EUROPEAN PARLIAMENT AND OF THE COUNCIL of 22 September 2010 on the protection of animals used for scientific purposes), as implemented in French legislation (February 1^st^ 2023), reviewed by local Ethics committee (CE71) and approved by French Ministry of higher education, research and innovation (project #1257).

Two male monkeys (*Macaca mulatta*, monkey R, ∼10 kg; monkey M, ∼9 kg) sat in a custom-made primate chair, head restrained, facing a 27-inch CRT monitor (resolution 2560*1440, 85Hz refresh rate, 42 cm viewing distance). Monkeys’ daily water supply was carefully controlled. Stimuli consisted of elongated colored patches whose orientation was evenly spaced across 180°. All stimuli were generated using the LAB color space (1976 CIE L*a*b) and all colors were measured as isoluminant under experimental conditions using a luminance meter (Minolta).

Gaze position and pupil size were measured with an optical eye tracker (SR Research) at 1.0 kHz sample rate. Reward delivery, stimulus presentation, behavioral signals and task events were controlled using MonkeyLogic software (Asaad et al., 2013), running under MATLAB on a Windows-based PC.

### Task

Both animals were trained to perform a delayed conjunction matching task. Trials began when subject grasped a manual lever located inside their chair and maintained gaze for 200 to 300 ms within a virtual window of 2° radius surrounding fixation point. Each trial consisted of a sample stimulus presentation (450 ms, either to the left or right hemifield), followed by a delay period (500 ± 100 ms for monkey R; 700 ± 100 for monkey M) and a sequence of variable number of successive test stimuli (450 ms each, on sample’s location). Up to five equiprobable test stimuli were presented in 31 of 53 sessions for monkey R, whereas up to four equiprobable test stimuli were presented in the remaining 22/53 sessions for monkey R and in all 24 sessions for monkey M. During each test epoch, a distractor stimulus was presented simultaneously in the opposite hemifield. Test and distractor stimuli were defined by the conjunction of one of eight colors (evenly distributed along a circle in LAB color space-1976 CIE L*a*b) and one of eight orientations (evenly spaced across 180°). Sample stimuli were defined by the conjunction of one of two opposite colors from the test set (color 1: c1 or color 5: c5) and one of two orientations (orientation 1: o1=0°, and orientation 5: o5=90°). Sample stimuli could be presented at one of two locations (diametrically opposed locations relative to the fixation point). Monkeys were required to release the manual lever when a test stimulus matched sample’s color, orientation and location to receive a reward (water or juice depending on animals’ preferences). In addition to the four possible non-neutral sample combinations (o1c1, o1c5, o5c1, o5c5), sample stimuli could also be neutral, consisting of a neutral color (central to color picking circle) and a neutral orientation (represented by a dot). In these neutral trials, test stimuli never matched the sample in either orientation or color, and no manual response was required. Likewise, in catch non-neutral trials, no target was presented. In both neutral and catch trials, monkeys were rewarded for holding the lever until the end of the trial.

If monkey failed to respond to currently presented target stimulus, trial stopped without reward. Any gaze deviation outside the virtual fixation window, or any response to a non-matching stimulus (false alarm responses), immediately stopped trial. Intertrial intervals (ITI) lasted 1500 ms for monkey R and 1200 ms for monkey M. Additional penalties were added to ITIs after fixation breaks (800 ms for monkey R, 5000 ms for monkey M), false alarm (3000 ms for monkey M) and misses (5000 ms for monkey M).

Behavioral performances analyzed here were acquired during 53 electrophysiological sessions in monkey R and 24 sessions in monkey M.

### Analyses

#### Gaze distance to stimuli projection over virtual fixation window

We tested whether gaze was attracted toward test stimuli locations. We used only correct trials for this analysis. Since stimuli’s eccentricity can vary for each session, we first projected sample stimuli’s coordinate onto the virtual fixation window (2° away from fixation point) and second computed its distance to the averaged eye position over a 200 ms window following each test onset (different windows give similar results).

#### Pupil size

We computed variations of pupil size for neutral and non-neutral trials at each test epoch. We used only correct trials for this analysis. We normalized raw signal (recorded with *open-ephys* system along with other electrophysiology signals which are not analyzed here) by dividing each trial pupil-size time series by their respective baseline defined as the average pupil size in a window 300 ms around sample onset (100 ms prior to 200 ms after sample onset). Pupil size traces were averaged over a 200 ms window aligned 100 ms after each test onset (other timing give similar results) or after sample offset (during the delay) during neutral and non-neutral trials were compared through ANOVA analyses and Tukey HSD tests.

#### Micro saccade rates

We computed micro-saccade rates (micro-saccades/second) at each epoch of the task (sample + delay, and each test epoch). We first smoothed each trial eye positions over time using averaging window of 50 ms moving every millisecond during each epoch of the task. Second, we computed the instantaneous eye acceleration (second derivative of eye position) at each time point of each trial. Micro-saccades were identified by detecting time points at which gaze acceleration exceeded monkey specific thresholds (2 and 2.5 standard deviations for monkey R and M, respectively). Accuracy of the micro-saccade detector was extensively assessed manually. Statistical comparisons between neutral and non-neutral trials at each epoch were computed using paired t-test (Bonferroni corrected). Comparison between task epochs was assessed using ANOVA analyses and Tukey HSD tests.

#### Decoding trial’s neutral status based on eye position and pupil size

We tested whether cognitive state (linked to sample’s neutral status) was reflected in eye position and pupil size. We trained a separate Support Vector Machine (SVM) classifiers for each session to decode neutral status based on eye position (cartesian coordinates centered on fixation point) alone or in combinations with normalized pupil size. We randomly picked 90 correct trials (45 neutral and 45 non-neutral trials) and sampled eye position and pupil size every 10 ms along trials (fixation period, sample presentation, delay period and succession of test stimuli). We tested decoder performances on a held-out set of 30 randomly picked correct trials (15 neutral, 15 non-neutral). These procedures were repeated 1,000 times, and classifiers were considered to perform above chance (50%) if the accuracy of the decoder was higher than chance level for more than 990/1,000 iterations (p < 0.01).

#### Decoding test epochs based on eye position and pupil size

We used a similar approach for decoding trials test epoch based on eye position and pupil size, except data were averaged on a 200 ms window aligned 50 ms after each test onset.

## Acknowledgement

We thank L. Condro for partial animal training, the staff of the Mediterranean Primate Research Center for expert veterinary assistance, the staff from the internal surgical platform (SurFiN) with a special emphasis on L. Renaud. We also thank F. Cazettes, G. Lindsay and members of the Invibe team for comments and discussion on earlier versions of this manuscript. *This work received support from the* European Research Commission Marie Sklodowska Curie RI program, ANR JCJC program, and the french government under the France 2030 investment plan, as part of the Initiative d’Excellence d’Aix-Marseille Université – A*MIDEX ” AMX-22-RE-AB-159. C.L was funded by Aix-Marseille University Neuroschool Phd Program. C.L. curated data and wrote custom code for data handling; A.M.A, A.F. and G.I. trained animals and acquired data; C.L. and G.I. analyzed the data; C.L. and A.F. assisted G.I. in writing the manuscript; G.I. conceptualized the study, designed the task and supervised A.M.A, C.L and A.F.

